# Modelling firing regularity in the ventral cochlear nucleus: mechanisms, and effects of stimulus level and synaptopathy

**DOI:** 10.1101/121707

**Authors:** Dan F. M. Goodman, Ian M. Winter, Agnès C. Léger, Alain de Cheveigné, Christian Lorenzi

**Affiliations:** Department of Electrical and Electronic Engineering, Imperial College London, UK; Department of Physiology, University of Cambridge, UK; Laboratoire des Systèmes Perceptifs, UMR CNRS 8248, Département d’études cognitives, Ecole normale supérieure, Paris Science et Lettres Research University, 75005 Paris, France; Department of Electrical and Electronic Engineering, Imperial College London, Exhibition Road, London SW7 2AZ, UK

**Keywords:** ventral cochlear nucleus, chopper cell, computational model, regularity, deafferentation

## Abstract

The auditory system processes temporal information at multiple scales, and disruptions to this temporal processing may lead to deficits in auditory tasks such as detecting and discriminating sounds in a noisy environment. Here, a modelling approach is used to study the temporal regularity of firing by chopper cells in the ventral cochlear nucleus, in both the normal and impaired auditory system. Chopper cells, which have a strikingly regular firing response, divide into two classes, sustained and transient, based on the time course of this regularity. Several hypotheses have been proposed to explain the behaviour of chopper cells, and the difference between sustained and transient cells in particular. However, there is no conclusive evidence so far. Here, a reduced mathematical model is developed and used to compare and test a wide range of hypotheses with a limited number of parameters. Simulation results show a continuum of cell types and behaviours: chopper-like behaviour arises for a wide range of parameters, suggesting that multiple mechanisms may underlie this behaviour. The model accounts for systematic trends in regularity as a function of stimulus level that have previously only been reported anecdotally. Finally, the model is used to predict the effects of a reduction in the number of auditory nerve fibres (deafferentation due to, for example, cochlear synaptopathy). An interactive version of this paper in which all the model parameters can be changed is available online.

**Highlights:**

- A low parameter model reproduces chopper cell firing regularity
- Multiple factors can account for sustained vs transient chopper cell response
- The model explains stimulus level dependence of firing regularity
- The model predicts chopper cells fire more irregularly after deafferentation
- An interactive version of the paper allows readers to change parameters

## 1 Introduction

The auditory system makes use of acoustic information on a wide range of temporal scales, from the microsecond scale of interaural time differences, up to the second or more scale of adaptation to stimulus statistics (Joris et al., 2004; McDermott and Simoncelli, 2011; McDermott et al., 2013). An important stage of temporal processing in the auditory system is the cochlear nucleus (e.g. Rhode and Greenberg 1992). Many neurons in this nucleus receive spikes directly from auditory nerve fibres (in addition to other structures), and these spikes retain stimulus-induced precise timing. Given this, the cochlear nucleus is expected to be an important neural structure for the initial extraction of temporal features to be used by the rest of the auditory system. Indeed, many cell types in this area show interesting temporal responses to stimuli, such as cells that respond only or primarily to the onset of a stimulus. All signals pass through the cochlear nucleus, so understanding how temporal structure is transmitted or processed by neurons in this nucleus is key to understanding processing in the whole auditory system.

A striking example of temporal structure in the spike trains of cochlear nucleus neurons is found in the *chopper* cell of the ventral cochlear nucleus, which initially responds to a pure tone stimulus with a highly regular spike train. Chopper cells have been suggested to be important in the coding of temporal envelope cues (e.g. Lorenzi et al. 1995; Carney et al. 2015; for a review see Joris et al. 2004), which is widely agreed to be essential for understanding speech (Shannon et al., 1995; Friesen et al., 2001). In some chopper cells, the initial regular firing is sustained over the whole duration of the tone, in which case the cell is classified as a *sustained* chopper. In other cases, this regularity is reduced after a few tens of milliseconds, in which case they are classified as *transient* choppers. This distinction is illustrated in Figure 1. A number of hypotheses have been put forward to explain this difference, including the effect of adaptation, the low-pass effect of dendritic filtering, the number of innervating fibres, and the degree of inhibition (Molnar and Pfeiffer, 1968; Blackburn and Sachs, 1989; Banks and Sachs, 1991; Hewitt et al., 1992; Hewitt and Meddis, 1993).

**Figure 1:**
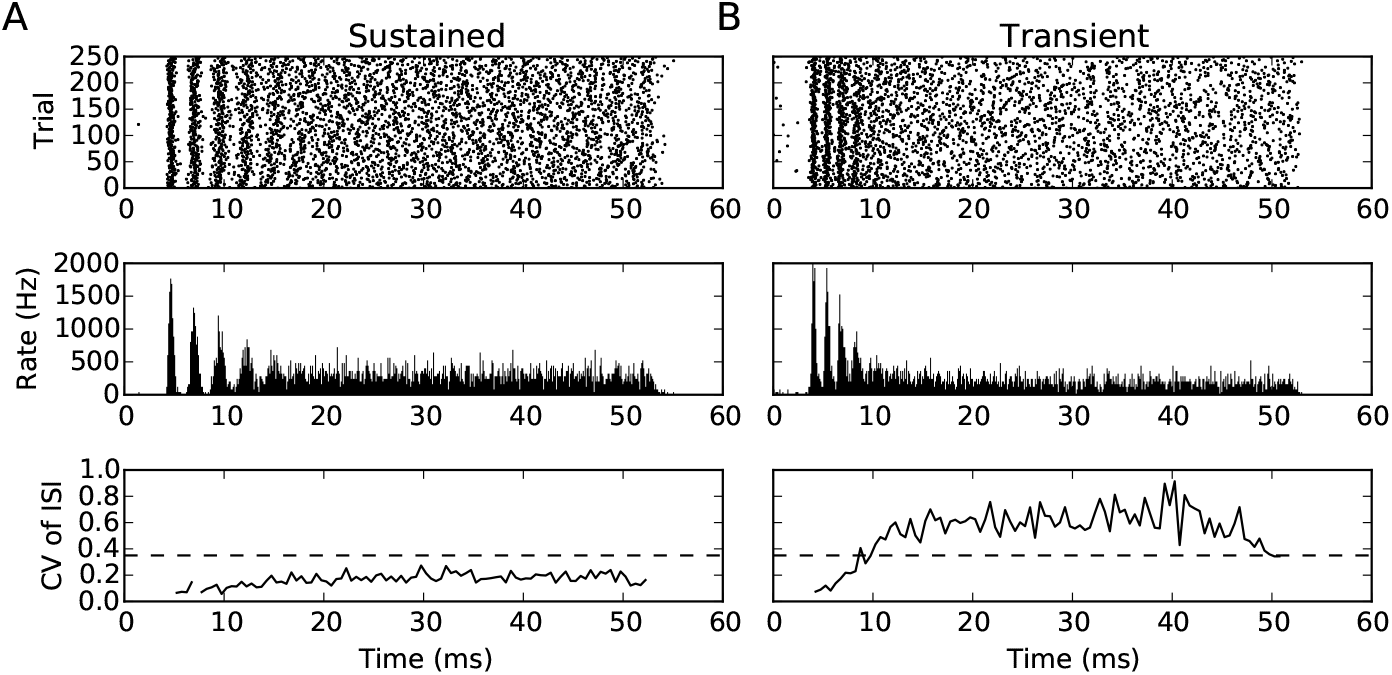
(A) Sustained and (B) transient chopper cells (recorded from the cochlear nucleus of the guinea pig). The upper panel shows the raster plot of the spikes in response to a tone at the BF of the neuron. The middle plot shows the instantaneous firing rate: note the initial regular firing over the first 10-15 ms of the tone. The lower plot shows the CV of the ISI distribution, with the dashed line showing the cutoff between sustained and transient behaviour. Note that both the sustained and transient cells show a flat histogram in the latter part of the stimulus, and so the shape of the histogram is not sufficient to distinguish them. This will happen whenever CV > 0 because the jitter in timing builds up at each spike.

So far, no consensus has been reached to explain the mechanisms underlying the difference between transient and sustained choppers. This is partly due to a lack of knowledge of key properties of these cells. Choppers are generally thought to be stellate cells, although the direct evidence for this is based on a relatively small number of cells for which anatomical and electrophysiological data have been collected (e.g. five sustained chopper cells in Smith and Rhode 1989). As an example of the problem, consider the number of auditory nerve fibres synapsing onto chopper cells. Most models assume a relatively large number (e.g. 60 in Hewitt and Meddis 1993), but experimental data on the number of inputs to stellate cells has a very wide range. Ferragamo et al. (1998) suggest a minimum as low as 5 auditory nerve fibre inputs, while Young and Sachs (2008) suggest at least 10-20, and note that the methods used in both papers may only be finding the strongest synapses, so that both may be substantial underestimates. Further complicating matters, recent physiological evidence suggests that there may not be clearly defined classes, but rather a continuum of cell types (Typlt et al., 2012; Rothman and Manis, 2003a,b).

Most models of chopper cells make rather specific assumptions about these parameters (Banks and Sachs, 1991; Guerin et al., 2006; Hewitt et al., 1992; Hewitt and Meddis, 1993; Rothman and Manis, 2003a, b; Wang and Sachs, 1995; Wiegrebe and Meddis, 2004). Different assumptions for different models make it difficult to compare them and determine what are the essential features of the models that contribute to their behaviour. This study presents a reduced mathematical model that reproduces the full range of observed response patterns on the basis of a small set of parameters, that can then be related systematically to observed or hypothesized anatomical and physiological mechanisms. The low number of parameters allows us to see which of these mechanisms have a common effect on the model variables and therefore response patterns. It also allows us to perform a thorough search of the parameter space to assess the robustness of the model (how much the behaviour depends on fine tuning of parameters), and to identify which variables – and therefore mechanisms – are critical to reproduce the observed patterns. For a further discussion of the advantages of this approach to modelling, see O’Leary et al. (2015).

In recent years, many researchers have investigated the idea of a “hidden hearing loss” (e.g. Kujawa and Liberman 2009; Schaette and McAlpine 2011; for reviews see Plack et al. 2014, 2016; Oxenham 2016; Liberman et al. 2016) in which neural output from the cochlea is reduced due to synaptopathy, but absolute (audiometric) thresholds may be normal or near-normal (so that the deficit is not detected in a standard clinical test). Kujawa and Liberman (2009) found that after an acoustic overexposure in which absolute thresholds only shifted temporarily, there was permanent damage to the cochlea. In particular, synapses were lost within 24h and auditory nerve fibres over several months. Schaette and McAlpine (2011) proposed (as a hypothesis to explain tinnitus) that the brain may compensate for this loss of input fibres with a homeostatic mechanism that increases gain centrally, boosting the synaptic strength of the remaining auditory nerve fibres in order to restore the original firing rates of neurons in the cochlear nucleus. In this study, we investigate the consequences of this hypothesis on the temporal response properties of cochlear nucleus neurons. We focus on temporal properties for two reasons. Firstly, by construction, cochlear nucleus neural firing rates will be left unchanged according to this hypothesis. Secondly, the cochlear nucleus is known to have an important role in the initial processing and transmission of temporal information to the rest of the auditory system.

The code for this paper is available online at https://github.com/neural-reckoning/vcn_regularity. It is also possible to interactively explore the figures in this paper online without having to install the software locally, including changing all parameters.

## 2 Methods

### 2.1 Chopper cells

Sound entering the auditory system is initially transduced in the cochlea leading to the firing of spikes in auditory nerve (AN) fibres. These auditory nerve fibres project excitatory connections to the cochlear nucleus (CN) which is divided into two areas, dorsal and ventral (DCN and VCN). Neurons in the CN receive both excitatory and inhibitory inputs (Campagnola and Manis, 2014). “Chopper” cells are located in the VCN, and are characterised by their initial very regular response to a tone at their best frequency (see figure 1). Choppers are further subdivided into sustained and transient categories. Sustained choppers continue to fire regularly throughout the duration of the tone, whereas transient choppers become more irregular over time (Pfeiffer, 1966; Rhode and Smith, 1986; Blackburn and Sachs, 1989; Winter and Palmer, 1990).

This regularity is measured using the coefficient of variation (CV) of the interspike intervals (ISIs): the ratio of the standard deviation to the mean. In the case of perfectly regular firing the standard deviation will be zero, so CV = 0. Poisson distributed spikes will have an ISI distribution with the mean equal to the standard deviation, and therefore CV = 1. The cutoff point between sustained and transient behaviours varies slightly in the literature, e.g. Young et al. (1988) use a cutoff of 0.35, while Blackburn and Sachs (1989) use 0.3. In this study, chopper responses are classified as sustained if the ongoing CV < 0.35, and as transient if CV > 0.35. Some examples of sustained and transient choppers are shown in figure 1.

### 2.2 Reduced model

Cells in the VCN have intricate morphologies, receive excitation and inhibition at different points on their dendritic trees and somas, and have complex cell properties with variable distributions of different types of ion channels. Several models that incorporate this level of detail have been constructed (e.g. Banks and Sachs 1991; Hewitt et al. 1992; Hewitt and Meddis 1993; Rothman and Manis 2003a, b). A strong driving force for the complexity of these models has been the desire to account for the catalogue of known properties of auditory neuron responses, including filtering, compression, adaptation, etc. A different approach is taken in the present study. Similarly to Molnar and Pfeiffer (1968), it is assumed that it is not necessary to reproduce all these known features in order to address a specific question. As Molnar and Pfeiffer were studying the *spontaneous* firing properties of VCN cells, they did not need to account for stimulus-specific response properties, including filtering, compression, and adaptation. They could therefore assume that the system was in a relatively stationary state. In other words, any one time interval was the same as any other. This led these authors to develop a very simple and yet effective model.

We will also assume that the system is in a partially stationary state, but in our case it is in response to a stimulus with a relatively long duration. Specifically, after a certain period of time we assume that all the slow adaptation processes have reached a stationary state, but not the rapidly varying membrane potential and post-spike refractoriness processes. Our model will therefore be able to make predictions about the ongoing but not the initial response to a stimulus. This is, however, sufficient for understanding the differences in ongoing firing regularity between sustained and transient choppers. In addition, we consider only pure tone responses at the cell’s best frequency; so, aspects related to cochlear filtering will not be taken into account. For most of the current study, we will also ignore the effect of stimulus level, so the effects of cochlear compression can be ignored. We will investigate level-dependence later in this study, but it will turn out that a detailed model of compression is not required to do so.

It is assumed that:

1. The cell is in a stationary state except for membrane potential and refractoriness dynamics.
2. Cells receive excitatory and inhibitory input.
3. The total incoming spike rate is high.
4. Incoming spikes follow a Poisson distribution with a fixed rate.
5. There are no noise correlations between the inputs.

The first assumption has been discussed above, and the next two are relatively uncontroversial. However the fourth and fifth may be violated. For example, the inhibitory inputs may violate these assumptions. One class of models of chopper cells includes inhibitory inputs driven partly by the same excitatory inputs as those that drive the chopper cell (e.g. Pressnitzer et al. 2001). This is not a possibility covered by the model presented here.

The first step in our model is to reduce the complexity of the neuron itself. Meng et al. (2012) showed that the widely used, detailed VCN model of Rothman and Manis (2003c) could be reduced to a much simpler two variable leaky integrate-and-fire (LIF) neuron. We go further and reduce to a one variable LIF neuron, which we can do as we have assumed that the cell is in the semi-stationary state described above and so the second adaptation variable is a constant. Specifically, we model chopper cells as leaky integrate-and-fire neurons with the following dynamics. The membrane potential *v* evolves continuously according to the differential equation:

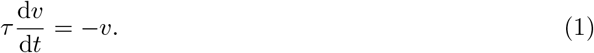

The constant *τ* is the effective membrane time constant. This can depend on the synapses and overall activity level, and may differ from the passive membrane time constant, which is why we refer to it as an effective time constant (Koch et al., 1996). An incoming excitatory (inhibitory) spike causes an instantaneous change *v* ← *v* + *w_E_*(*v ← v – w_I_*). We refer to *w_E_* (*w_I_*) as the excitatory (inhibitory) synaptic strength. The neuron fires a spike when the membrane potential crosses the threshold *v* > 1 and immediately resets to *v* ← 0. The neuron then stays refractory with *v* = 0 for a time *t*_ref_.

This is a highly simplified model of a neuron, but captures all the important dynamics if we assume that it has reached the semi-stationary state described above. The effect of slow adaptation processes are not ignored in this model, but are reflected in the parameters *w_E_* and *w_I_*, as they are equivalent to a simple scaling. As we do not consider the full range of fast dynamics, we could not, for example, model a bursting cell, but this is not necessary for the cells under consideration. We ignore very fast synaptic dynamics, modelling them as being instantaneous. Synaptic dynamics can be easily incorporated into the model, and do not substantially change the results (data not shown). Finally, for the neuron’s morphology, we can capture the effect of low pass dendritic filtering by simply increasing the effective time constant *τ* (Sumner et al., 2009).

### 2.3 Diffusion approximation

With the additional assumptions that: *w_E_ = w_I_ = w*; there are *N* excitatory and *N* inhibitory incoming synapses; each excitatory (inhibitory) input fires Poisson spikes at a fixed rate *ρ_E_* (*ρ_I_*); and that *Nρ_E_* and *Nρ_I_* are relatively large, we can further simplify this model using the *diffusion approximation*. Fourcaud and Brunel (2002) showed that this model can be well approximated by the stochastic differential equation

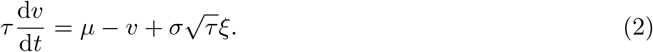

The term *ξ* is a stochastic differential, *μ* and *σ* are constants, and the threshold and resetting behaviour are as before. The stochastic differential ξ can be thought of, over a time window [*t, t + δt*] of width *δt*, as a Gaussian random variable with mean 0 and variance 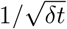. The constant *μ* reflects the mean combined excitatory and inhibitory current, and *σ* reflects the variation in that current. The constants *μ* and *σ* can be computed as

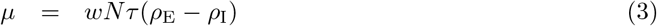

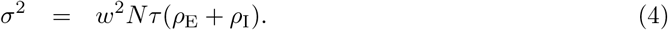

Ultimately, the behaviour of this approximation depends only on the values of *μ, σ, τ* and *t*_ref_. It depends on *N, w, ρE* and *ρ_I_* only insofar as *μ* and *σ* depend on these. So, while the behaviour of the model nominally depends on the assumption of *N* excitatory and *N* inhibitory inputs with the same synaptic weights, a different choice will only change the way we compute *μ* and *σ* in equations 3 and 4, but not the actual dynamics of the model specified in equation 2. For this reason, in section 3.2 we will study the behaviour of equation 2 as we vary *μ* and *σ* directly (rather than computing them from *N, w, ρ_E_* and *ρ_I_*), and these results will apply more generally than when we make a specific assumption about the number of excitatory and inhibitory inputs.

Figure 2 illustrates that the diffusion approximation reproduces well the time course of the membrane potential. Although the form of equation 2 is more complex than that of equation 1, overall the model is considerably simpler. Whereas previously we had 2*N* + 1 neurons in our model (including the *N* excitatory and *N* inhibitory inputs), we now only have one. In addition, the complex effect of these synapses is captured in a single equation allowing us to understand the behaviour more clearly. We will make use of this in section 3 (for example, in equation 5).

**Figure 2:**
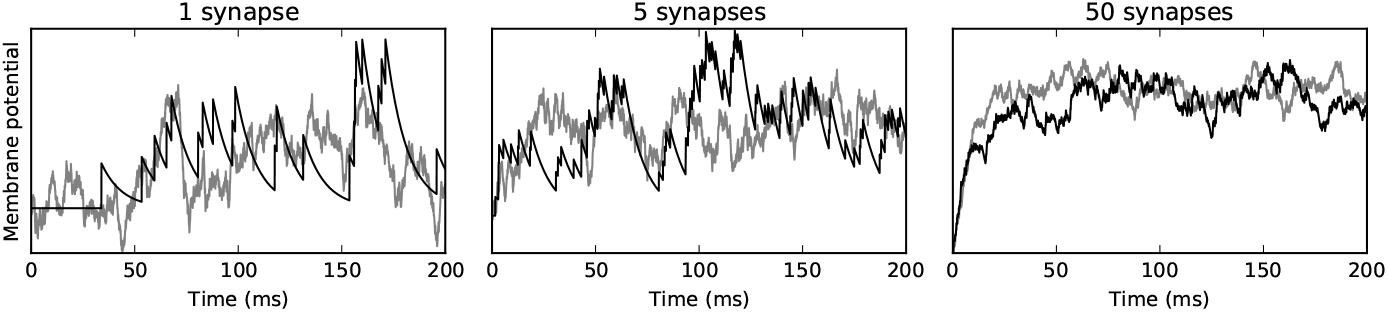
Diffusion approximation. In each panel, the black curve shows the behaviour of the model neuron, and the grey curve the diffusion approximation. Note that the processes are random, so we do not expect the approximation to match exactly, only in the overall behaviour (size and timing of excursions). The panels show the approximation for 1, 5 or 50 input synapses. For 5 or more synapses, it is difficult to distinguish which is the model and which is the approximation, and even for 1 synapse, the size of the larger excursions matches, and only the small scale structure differs.

It is possible to find an analytical solution for the firing rate and CV of this model. However, we opted to use simulated results rather than the analytical solution for a number of reasons. Firstly, it is not possible to find a closed form solution. Secondly, the solution is specific to this precise equation and not straightforward to adapt to other similar models. Thirdly, computing the integrals below numerically is not straightforward, especially when σ is small, and is actually slower than running the simulation.

The following is derived from Brunel (2000). Without refractoriness, the mean interspike interval is

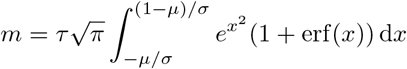

and so the firing rate is 1/*m*. The CV is

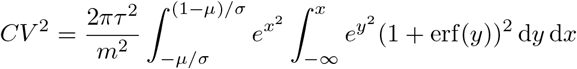

With refractoriness, the mean interspike interval is 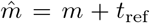, with corresponding firing rate 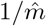. The CV is 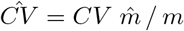.

### 2.4 Level dependence

In section 3.3 we will describe how the model can account for responses to tones at different sound levels. We model this by choosing two sets of parameters, one at the 20 dB level, and one at the 50 dB level (all levels are given re threshold at best frequency in this section). We describe the method for choosing these parameters below.

In figure 7 we select parameters uniformly randomly and reject certain parameters according to the procedure described below. With this random distribution of parameters we show the density of the joint distribution of chopper cell firing rates and CVs. This density plot is computed by generating 10,000 sets of parameters, and then smoothing the data using a Gaussian kernel density estimation. This is implemented using the `gaussian_kde` method in the scipy Python package (Jones et al., 2001) with Scott’s rule for the bandwidth selection method (Scott, 1992). The parameters are chosen as follows. The following parameters are the same at both stimulus levels: the number of excitatory and inhibitory input neurons *N* is chosen uniformly between 3 and 100; the membrane time constant *τ* between 1 ms and 10 ms; the refractory period *t*_ref_ between 0 ms and 1 ms. The mean current *μ_20_* at the 20 dB level is chosen uniformly randomly between 1 and 5. This choice of μ_20_is used to calculate the synaptic weights *w*, which is then unchanged at the 50 dB level allowing us to compute the mean current *μ_50_* at that level. The excitatory firing rate and fraction of inhibition at the 20 dB level, *ρ*_20_ and *α*_20_ are chosen between 100-300 sp/s and between 0 and 0.8 respectively. The change in these two parameters as we increase the level from 20 dB to 50 dB is: *ρ*_50_ – *ρ*_20_ between 0 and 160 sp/s; *α*_50_ – *α*_20_ between −0.3 and 0.5. Finally, if this leads to a chopper firing rate outside of the range 100 to 500 sp/s, or a CV outside the range 0.1 to 0.7, then it is rejected (as this is the range we observed experimentally).

In figure 8 we also select parameters randomly, but in this case we use a choice of distributions designed to mimic the experimental data more closely. The mean level of excitation at the 20 dB level *μ*_20_is chosen as a normal random variable with mean 2 and standard deviation 0.4. The fraction of inhibition *α* at each level is chosen independently as 0.65*f (x)* where *x* is uniformly distributed between 0 and 1, and *f (x)* = 1/(1 + *e*^6(1-2*x*)^). This allows for the amount of inhibition to go up, down, or stay the same at the 50 dB level compared to the 20 dB level. The excitatory firing rates *ρ*_E_ = *ρ* are chosen for each level as a normal random variable with mean 250 sp/s and standard deviation 25 sp/s. The number of excitatory and inhibitory inputs *N* is chosen as a uniformly random integer between 30 and 60. The log time constant (in ms) log *τ* is chosen uniformly between log5 and log15, and log refractoriness (in ms) log *t*_ref_ uniformly between log 0.1 and log 5. Finally, if one of the following conditions is violated then a new random selection of all the parameters is made: 1 < *μ* < 4; 150 < *ρ* < 450; the excitatory firing rate at 50 dB is higher than the excitatory firing rate at 20 dB; the difference between the inhibitory fractions α at the two levels is between −0.1 and 0.25. This distribution may seem complicated, but the idea is simple: most parameters are uniformly, normally or exponentially distributed in a reasonable range to roughly match the data, and the inhibitory fractions are chosen to have a bimodal distribution to reproduce the bimodality in the distribution of the CVs in the data.

### 2.5 Deafferentation

In section 3.4 we will show how a loss of auditory nerve fibre inputs (deafferentation) affects chopper cell firing regularity. We model this by reducing the number of auditory nerve fibre inputs *N* and increasing the synaptic weights w to restore the original chopper cell firing rate. We compute this by varying *w* until we find a firing rate that is close to the original, and then once a sufficiently large number of pairs of synaptic weights with their associated firing rates have been found, we smooth and interpolate these to plot them.

### 2.6 Amplitude modulation

In section 3.4 we will show how deafferentation may impact amplitude modulation tuning of chopper cells. We model amplitude modulation processing by assuming that the firing rate variable *ρ* is replaced by an amplitude modulated firing rate varying over time as

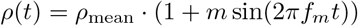

*ρ(t)* varies with a modulation frequency of *f_m_* with a mean value of *ρ_mean_* with a modulation depth of *m*. We model the effect of deafferentation followed by homeostatic plasticity by reducing the number of auditory nerve fibre inputs *N* and increasing the synaptic weights *w* to maintain the same mean input current *μ*, as before. In figure 10, the specific parameters used were: *ρ_mean_* = 200 sp/s; mean input current at *ρ = ρ*_mean_ was set to *μ*_mean_ = 1.25; *τ* = 10 ms; *t*_ref_ = 1 ms; *m* = 0.25. For the sustained chopper, inhibition was set to *α* = 0 and for the transient chopper *α* = 0.4. For the normal fibres, *N* = 50 and for the impaired fibres, *N* = 10.

In figure 10 we use the vector strength to measure the temporal modulation transfer function. This is computed in the standard way as the modulus of the mean modulation phase represented in the unit circle. 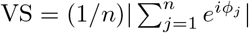, where *ϕ_j_* is the phase of spike *j* (of *n* spikes total) with respect to the modulation frequency.

Note that an amplitude modulated signal violates the assumptions of the model listed in section 2.2. We discuss this further in section 3.4.

### 2.7 Implementation

The model was simulated with the “Brian” spiking neural network simulator (Goodman and Brette, 2008, 2009; Stimberg et al., 2014). Each run was repeated 4000 times, for a duration of 350ms of simulated time per run. Spikes in the first 100ms were discarded, and the CV was computed as the ratio of the standard deviation to the mean of the simulated interspike intervals for the remaining 250ms.

The code is available at https://github.com/neural-reckoning/vcn_regularity.

### 2.8 Experimental methods

Experiments in the ventral cochlear nucleus were performed on anaesthetized pigmented guinea pigs (*Cavia porcellus*), weighing between 320 and 600 g. Detailed methods have been reported elsewhere (e.g. Ingham et al. 2016). Single units were recorded extracellularly with glass-coated tungsten microelectrodes. Upon isolation of a unit, its best frequency and excitatory threshold were determined using audio-visual criteria. Spontaneous activity was measured over a ten-second period. Single units were classified based on their peri-stimulus time histograms (PSTH), the first-order interspike-interval distribution and the coefficient of variation (CV) of the discharge regularity. The CV was calculated by averaging the ratios of the mean ISI and its standard deviation between 12 and 20 ms after onset (Young et al., 1988). PSTHs were generated from spike-times collected in response to 250 sweeps of a 50-ms tone at the units BF at 20- and 50-dB above threshold. Tones had their starting phase randomized and they were repeated with a 250-ms period. The experiments performed in this study have been carried out under the terms and conditions of the project license issued by the United Kingdom Home Office to the second author.

## 3 Results

### 3.1 Model behaviour

We start by giving some intuition for the behaviour of the model developed in section 2 and defined by equation 2 before presenting the results in detail. Recall that this is a model only of the ongoing and not the initial part of the response of a chopper cell to a pure tone stimulus. The model states that the membrane potential *v* fluctuates as an Ornstein-Uhlenbeck process (similar to a Brownian motion) around a mean value *μ*, with a time scale *τ*, and a spatial scale *σ* (so that the majority of the time it will be within *μ ± σ* but will occasionally fluctuate outside this range). Whenever it crosses the threshold *v* > 1 it fires a spike and resets instantaneously to *v* = 0 and stays there for the refractory period *t*_ref_. We illustrate the behaviour of the model for two key operating regimes in figure 3: the mean-driven and fluctuation-driven regimes (Shadlen and Newsome, 1998; Renart et al., 2007). In the mean-driven regime, the mean value *μ* > 1 is higher than the threshold for firing a spike, and the fluctuations *σ* are small, so the neuron will fire regularly. We measure this regularity with the coefficient of variation (CV) of the interspike intervals, which varies from 0 for a perfectly regular spike train to 1 for the highly irregular Poisson process (see section 2). The CV for the neuron in the mean-driven regime will therefore be low, and it will behave as a sustained chopper (CV< 0.35). In the fluctuation-driven regime, the mean value *μ* < 1 is below threshold and is therefore not sufficient to cause the neuron to fire a spike, but the large value of *σ* means that the random variations of *v* around the mean occasionally cause it to cross the threshold and fire a spike. In this case, the firing times will be much more random, the CV will be higher, and it will behave as a transient chopper (CV> 0.35).

**Figure 3:**
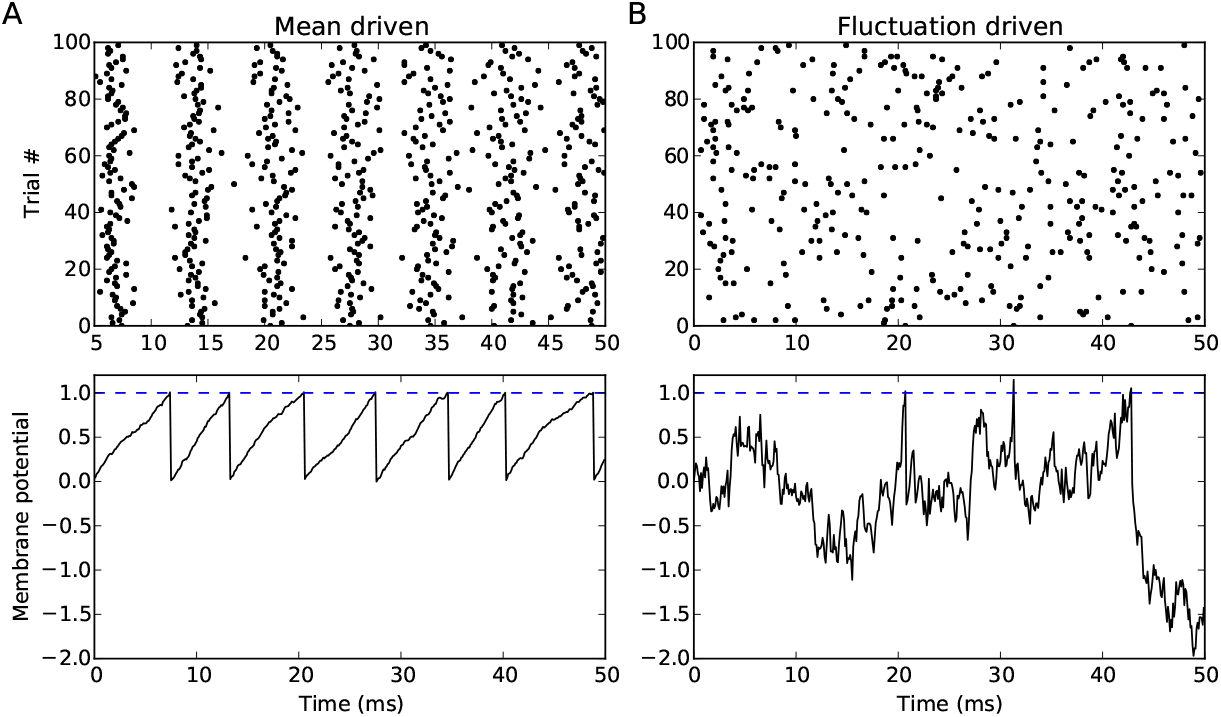
Mean-versus fluctuation-driven regimes of neural firing. Upper panels show raster plots over several repeats, lower plots show a single sample evolution of *v* over time, with the firing threshold indicated by the dashed line. (A) Mean-driven regime: the mean current *μ* > 1 and the noise level *σ* ≪ *μ* so the neuron is regularly driven to fire. The mean-driven regime corresponds to the behaviour of a sustained chopper. (B) Fluctuation-driven regime: the mean current *μ* < 1 and the noise level *σ* is large, so the neuron fires irregularly in response to random fluctuations in the current. The fluctuation-driven regime corresponds to the behaviour of a transient chopper.

Intuitively, if *σ/μ* is small, we would expect regular firing, and if *σ/μ* is large, we would expect irregular firing. Writing the firing rate of the inhibitory inputs *ρ_I_ = αρ*, and of the excitatory inputs as *ρ_E_ = ρ* and using equations 3 and 4 we can compute *μ* and *σ* as follows (recall from section 2 that *N* is the number of excitatory and inhibitory inputs):

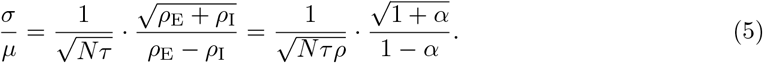

Two conclusions can be drawn from this. Firstly, if there is no inhibition (*α = ρ*_I_= 0) then the regularity will depend on the total incoming spike rate *N_ρ_*. The higher the rate, the more regular. Similarly, even if there is inhibition, the larger the number of incoming synapses, the more regular the output will be. Secondly, increasing inhibition will always make the output more irregular. An intuitive way to understand this last fact is as follows. If *X* and *Y* are two positive random variables, then the mean of *X – Y* is the difference of the means of *X* and *Y* (i.e. it is smaller than the mean of *X*), whereas the variance of *X – Y* is the sum of the variances of *X* and *Y* (i.e. it is larger than the variance of *X*).

### 3.2 Factors underlying chopper-like behaviour

Figure 4 shows how the firing rate and CV vary depending on the values of *μ* and *σ*. The precise shape of this map depends on the values of the parameters *τ* and *t*_ref_, however there are some common features. Typically, the larger the mean input μ, the higher the output firing rate. Moreover, as predicted, the larger the ratio *σ/μ* the higher the CV. This is not true in all cases: the shape of the map is more complex in regions where the output firing rate is low (where both *μ* and *σ/μ* are small). Understanding these areas would be important for understanding the spontaneous firing of the cell, or its behaviour near threshold, but typically firing rates are relatively high in the stimulus driven regimes that have been used to distinguish between sustained and transient choppers. Figure 4 shows the same plot for a few different values of *τ* and *t*_ref_. We do not show detailed plots of how CV depends on these two variables, as the dependence on *μ* and *σ* is the most important, but it can be summarised briefly by saying that larger values of either *τ* or *t*_ref_ tend to promote regularity.

**Figure 4:**
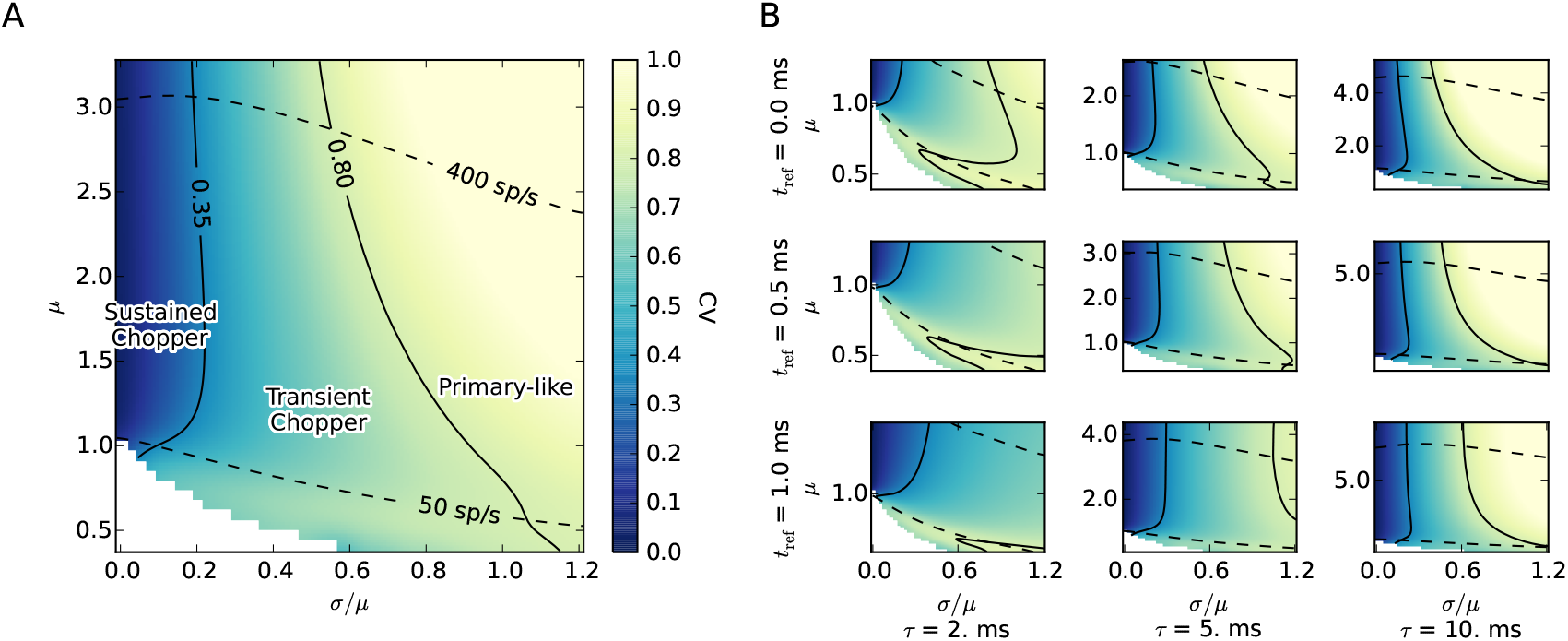
Map showing how firing rate and CV vary depending on *μ* and *σ*. (A) Detailed view for *τ* = 6 ms and *t*_ref_= 0.1 ms.Colour indicates the CV of the output spike train. The white areas show where there were no spikes, or insufficiently many to compute the CV. The solid black lines are contours indicating when CV is 0.35 and 0.8 (as none of the recorded chopper cells had a CV above this level, and it is more characteristic of primary-like cells). The dashed black lines are contours indicating when the output firing rate is 50 sp/s and 400 sp/s. (B) Same map but for different values of *τ* and *t*_ref_. Note that the (vertical) *μ* axis is different for each plot as the relationship between input spike rates and the value of *μ* depends on *τ* and *t*_ref_.

Also note that a region labelled “primary-like” has been included in this figure, as the model developed above is capable of reproducing the response of other cell types than chopper cells. In this case, we can see that when *σ/μ* is relatively large, the CV will be larger than 0.8 and this is more typical of a primary-like cell than a chopper cell. In principle the model also applies to cells such as onset cells, but since it only predicts the ongoing not the initial part of the response, it will not be very interesting in this case. Similarly, the current model cannot be applied to study the difference between primary-like and primary-like with notch, since the presence or otherwise of the notch is in the initial part of the response.

We can now use this model to explore the different factors that have been suggested to underlie chopper-like behaviour. The simplest of these is the number of inputs to the cell. Suppose we fix the mean input level *μ* and we wish to change the number of inputs *N*. From equation 3, we can do this by modifying the synaptic weight to be *w = w*_ref_/*N*. Now, from equation 5 we can compute that *σ/μ* is proportional to 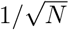. In other words, increasing (or decreasing) the number of inputs while leaving *μ* fixed will decrease (or increase) *σ/μ* (a horizontal translation in Figure 4), and therefore decrease (or increase) the CV. More inputs leads to more regular firing, while fewer inputs leads to more irregular firing. The number of inputs is therefore a plausible candidate mechanism to explain the difference between transient and sustained chopper cells. Transient choppers would have fewer inputs, while sustained choppers would have more.

We can carry out a similar analysis for other factors. Figure 5 shows the results of varying the number of inputs *N*; the degree of inhibition with respect to excitation *α*; the membrane time constant *τ*; and the excitatory input spike rate *ρ*_E_. Varying the number and strength of inhibitory inputs would lead to a modification of *α*. Varying the location of inputs from the soma to the dendrites would lead to a change in the effective membrane time constant *τ*. The effect of adaptation would be equivalent to a change in the synaptic weight *w* and therefore in the mean excitation *μ* (i.e. just a vertical translation in any of these maps). Varying any of these parameters is enough to introduce regions of sustained and transient chopper behaviour. Given natural variations in all these parameters in real chopper cells, we argue that it is unlikely that there is a single factor underlying the difference between sustained and transient chopper cells, but rather that there is a continuum of types of both sustained and transient chopper cells differing on one or more of these axes. We will return to this point in section 4.

**Figure 5:**
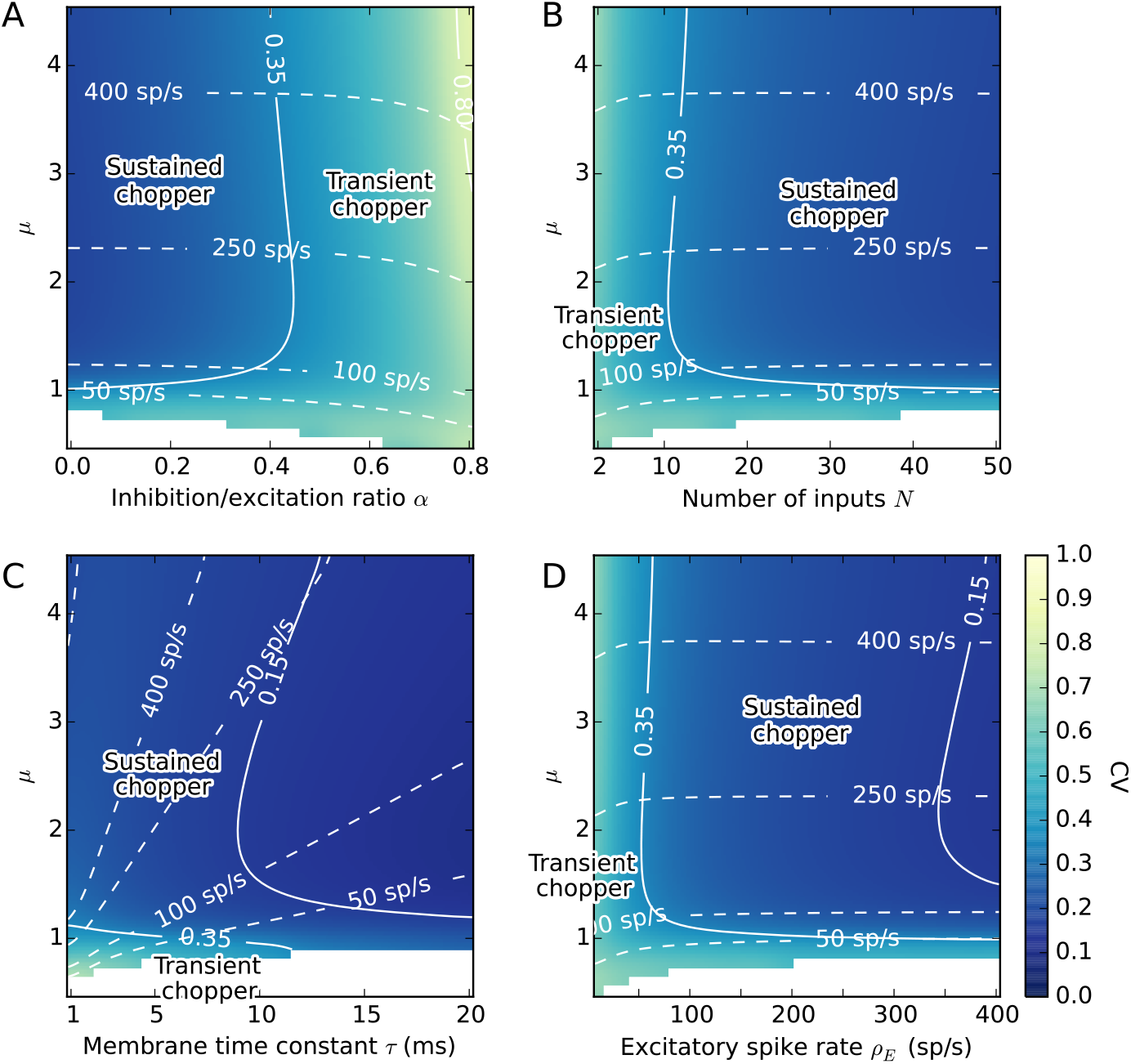
Map showing how CV and firing rate depend on five parameters of the model. In all cases, the mean excitation *μ* varies on the vertical axis, the colour indicates the CV, the dashed lines are firing rate contours and the solid lines are CV contours (as in Figure 4). The white areas show where there were no spikes, or insufficiently many to compute the CV. The horizontal axis varies (A) the ratio of inhibition to excitation *α*, (B) the number of inputs *N*, (C) the membrane time constant *τ*, (D) the excitatory firing rate *ρ*_E_. Each of these variables can account for both sustained and transient chopper cells.

### 3.3 Level dependence

Classification of chopper cells is carried out at a fixel level, or at a fixed level relative to threshold. However, based on the model above, we would expect CV to vary as a function of level. For example, at a higher level, the auditory nerve fibre inputs to the chopper cell would be firing at a higher rate (increasing *ρ*_E_). This would lead a change in both *μ* and *σ/μ*. We would therefore expect that not only would the chopper cell’s firing rate be level-dependent, but also its CV.

Figure 6 shows a new analysis of an existing data set where tones were presented at 20 dB re threshold and 50 dB re threshold. Almost all cells do indeed show a change in both firing rate and CV at these two different levels. In the majority of cases, a cell classified as sustained or transient at one level, would also be so classified at the other level. However, in 7% of cells (6 cells total), as the sound level moves from 20 dB to 50 dB re threshold, the CV increases from less than 0.35 to higher than 0.35, corresponding to a change in classification from sustained to transient. This pattern was also observed in a few cells by Blackburn and Sachs (1989), although they did not present general trends in changes in firing rate and CV for a population of cells.

**Figure 6:**
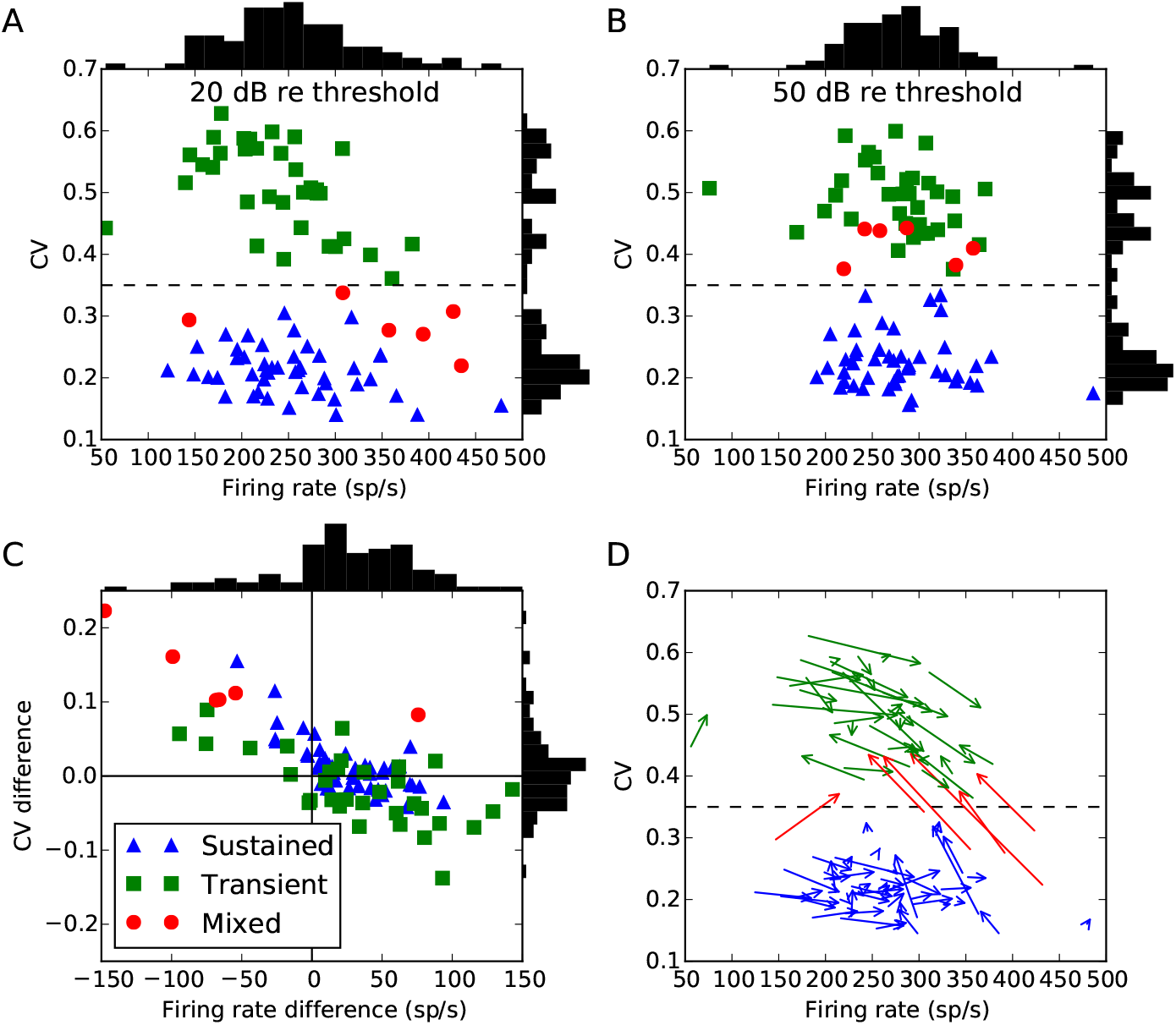
Chopper cells at 20 and 50 dB re threshold. Points labelled sustained (blue) or transient (green) would have been so classified at either level, whereas points labelled mixed (red, around 7% of all the cells) would have changed classification. Vertical histograms show distribution of firing rates, and horizontal histograms for CV. (A) CV and firing rate at 20 dB, and (B) at 50 dB. (C) Differences in firing rate and CV from 20 to 50 dB. The bottom right quadrant shows cells where the firing rate increases along with an increase in regularity, which can be explained simply by an increase in excitation. The upper left quadrant shows cells whose firing rate and regularity both decrease, which can be explained by an increase in inhibition. The upper right quadrant shows cells whose firing rate increases but fire more irregularly, which can be explained by an increase in both excitation and inhibition. There are no cells in the lower left quadrant, which would correspond to a lower firing rate but increased regularity, which would be difficult to explain as it would require either decreased excitation or increased inhibition, both of which lead to more irregular firing. (D) Arrows go from value at 20 dB to value at 50 dB.

The data shows a general negative trend between changes in firing rate and CV going from 20 dB to 50 dB re threshold. An increase in firing rate will tend to be associated with a decrease in CV, and a decrease in firing rate with an increase in CV. There are some cases where firing rate and CV both increase as level increases, but none in which both firing rate and CV decrease. Given that auditory nerve fibres have monotonically increasing rate-level curves, a decrease in firing rate at higher levels suggests an increase in inhibition. This is supported by the fact that adding inhibition always decreases regularity, and all cells in which firing rate decreased at higher levels also experienced a decrease in regularity. Similarly, an increase in the chopper cell firing rate is likely to be associated to an increase in the amount of excitation, which in the absence of inhibition will decrease *σ/μ* and therefore usually decrease the CV. The remaining cases in which both firing rate and CV increase may be caused, for example, by an increase in both excitation and inhibition (as a relatively small increase in inhibition can lead to a large increase in *σ/μ*).

The model allows each of these dependencies to be investigated in detail. However an exhaustive exploration of the four dimensional parameter space (*μ, σ, τ, t*_ref_) at the two different levels is hard to visualize and confront with available experimental data. For this reason, we approach the problem in two ways. For the printed paper, we plot the average behaviour over a random sampling of the parameters across a wide range. For the online, interactive version the full parameter space can be searched by selecting which parameters to fix and which to vary. In practice, the behaviour in general does not widely differ qualitatively from the average behaviour, suggesting that the results are robust.

Figure 7 shows how the firing rate and CV of the model chopper cell change as level increases from 20 to 50 dB over a wide range of parameters. We characterise the effect of a change in level purely as a change in the excitatory and inhibitory input firing rates. Excitatory rate is only allowed to increase (assuming that the primary excitatory drive is from the auditory nerve), whereas inhibition can either increase or decrease (since cells in the CN can have non-monotonic rate-level functions). Varying the two parameters *ρ*_E_ and the ratio of inhibition to excitation *α* in this way reproduces quite well the general negative trend between chopper cell firing rate and CV seen in the data, as well as the fact that firing rate and CV can occasionally both increase, but not both decrease. This trend is seen without needing to tune the parameters of the model, as changing the lower and upper limits of the parameter ranges does not substantially change the shape of this figure, as can be explored in the online interactive version.

**Figure 7:**
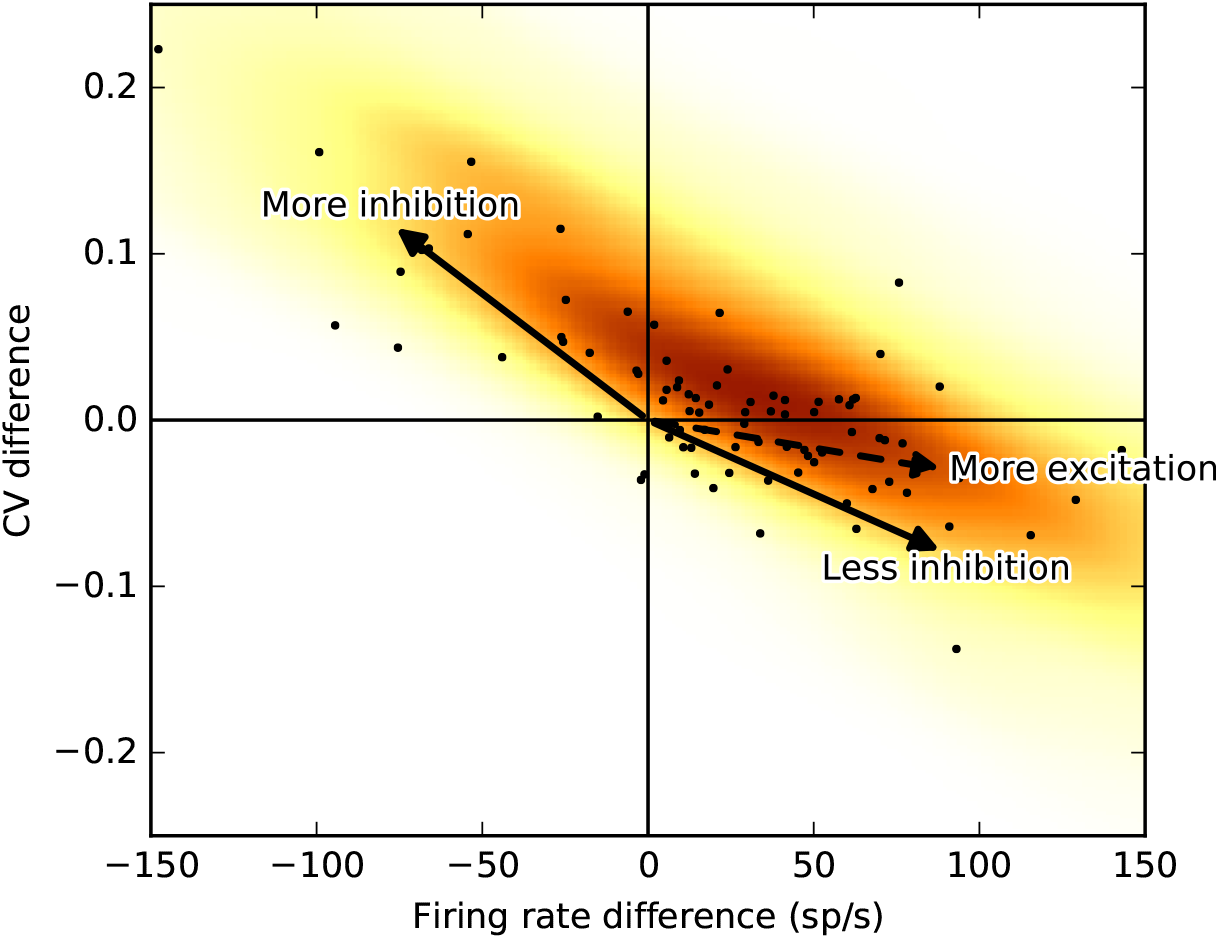
Level dependence of model chopper cell firing rate and regularity. Shows the change in chopper cell firing rate and CV as sound level increases from 20 to 50 dB re threshold. Black points show experimental data (as in Figure 6C). The colour of each point shows how frequently that rate and CV difference occurs in the model with uniformly randomly selected parameters (darker coloured areas occur more frequently).

Figure 8 shows the results of a model based on a random selection of the parameters *μ, ρ, α, N, τ* and *t*_ref_, where the distributions are chosen to more closely reflect the data. The precise details are given in section 2, but in short each parameter is chosen according to a uniform or normal distribution, except for the degree of inhibition which is chosen with a bimodal distribution to reproduce the observed bimodality of the CVs. Parameters that lead to chopper cell firing rates that are outside the observed ranges are rejected. Since the details of the distributions of the parameters are arbitrarily selected, we do not claim that this is a faithful model of real populations of chopper cells. The point is simply to demonstrate that the model can reproduce the full range of response patterns found in the experimental data in Figure 6. We used inhibition to account for bimodality of the CV distribution, but other choices would have worked as well.

**Figure 8:**
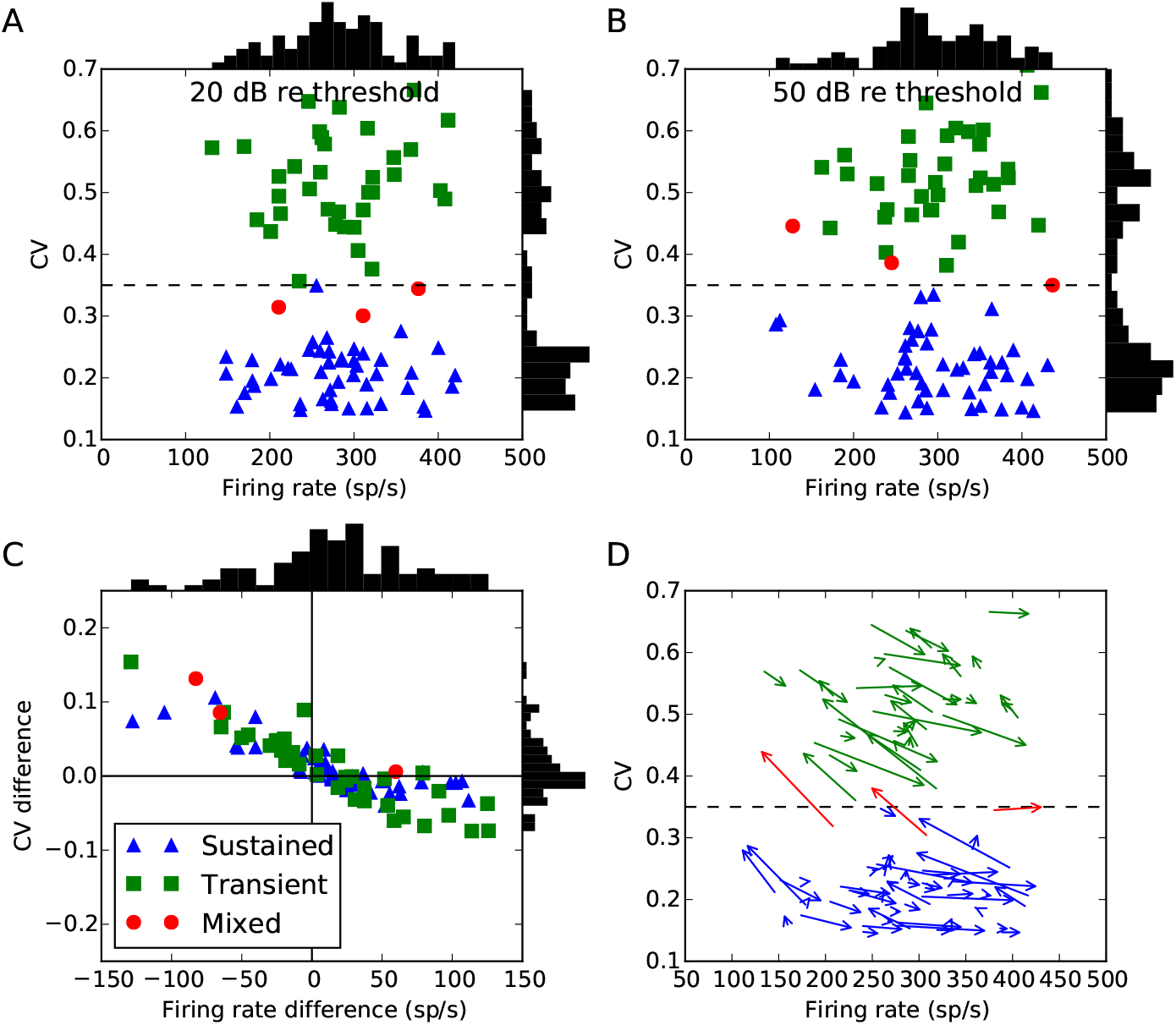
Model prediction of how chopper firing rate and CV change. Compare to experimental data in Figure 6.

### 3.4 Deafferentation

As a final result, we predict how the behaviour of these cells would change as a function of deafferentation, or loss of auditory nerve fibres. We will assume that after deafferentation, initially there will be a decrease in chopper cell firing rates reflecting a decrease in the total amount of excitation received from the auditory nerve (as some of the fibres will be lost). However, we assume a homeostatic mechanism increases the synaptic weights to compensate for this and restore the original chopper firing rate (Turrigiano, 1999, 2012). There is evidence that homeostatic changes occur in multiple regions of the auditory system from the cochlear nucleus to cortex, in response to various types of damage (Brozoski et al., 2002; Schaette and McAlpine, 2011; Gu et al., 2012; Chambers et al., 2016; Shore et al., 2016; Asokan et al., 2017) We assume that the only homeostatic changes are to the synaptic strengths and not to the intrinsic excitability of the cell as the intrinsic properties of the cell do not change substantially after complete deafferentation (Francis and Manis, 2000).

We can get an initial sense of the effect of deafferentation using the calculation from an earlier section on the effect of changing the number of inputs *N*. We saw that if we fix *μ* by setting the synaptic weight *w = w*_ref_/*N* (an approximation to a homeostatic process that restores the output firing rate), then *σ/μ* is proportional to 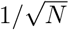. In other words, decreasing *N* (deafferentation) will lead to an increase in *σ/μ*. Since CV in general increases with *σ/μ*, deafferentation will in all cases lead to an increase in CV, and consequently a decrease in firing regularity.

Figure 9 shows a concrete example of this. In this figure, we assume only excitatory inputs but similar results would apply if we included inhibition. We generate this figure by simply selecting the number of inputs *N* and the desired output firing rate, and then varying the synaptic weight w until this firing rate is achieved. Deafferentation corresponds to a decrease in the number of inputs *N* (i.e. a move to the left in the figure) along with an initial decrease in the firing rate (downwards move), followed by the homeostatic restoration of the original firing rate (upwards move to the same original height). For this figure then, the long term effect of deafferentation corresponds simply to a jump to the left, which in almost all cases leads to more irregular firing (increased CV) although can in a very few cases lead to no effect or the opposite effect for some values of the parameters.

**Figure 9:**
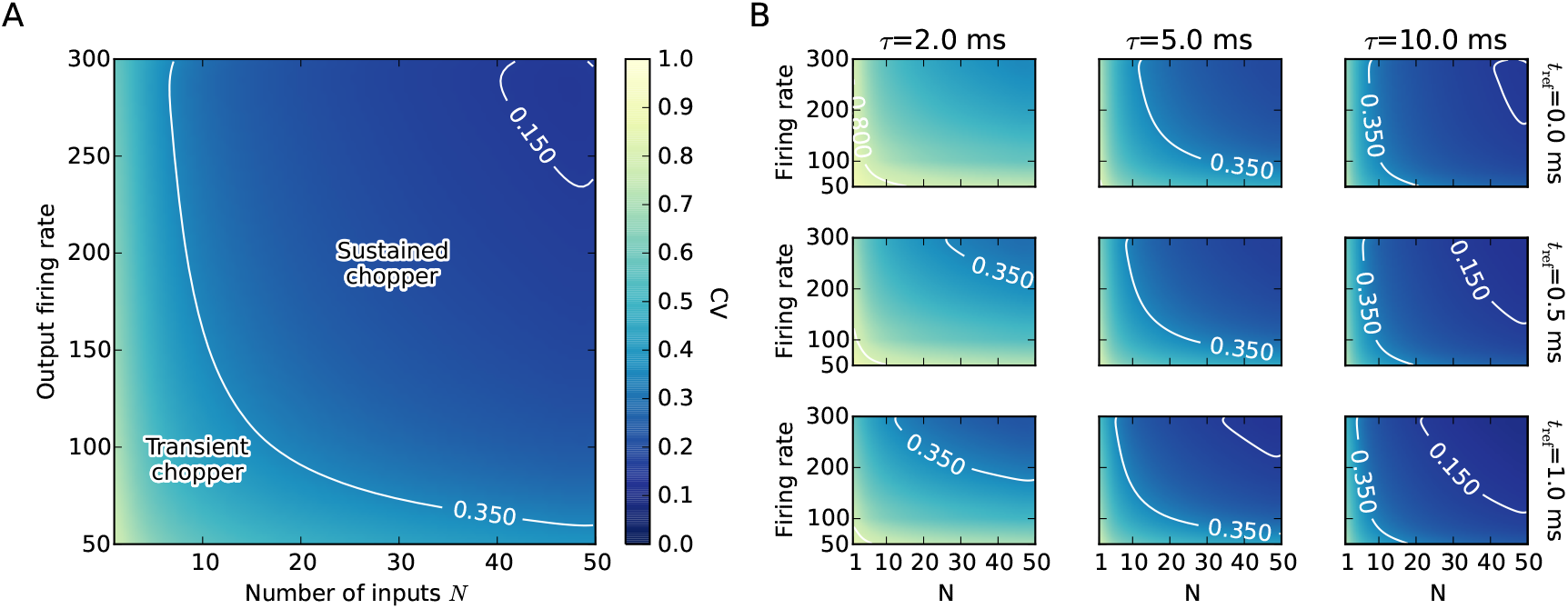
Map of CV as a function of the number of auditory nerve inputs and the chopper cell firing rate. The long term effect of deafferentation is a translation to the left. (A) Detailed view for *τ* = 6 ms, *t*_ref_ = 0.6 ms. (B) Same map for different values of *τ* and *t*_ref_. White lines are CV contours at 0.35 and 0.15. The latter contour is shown because very few of the recorded cells had a CV below this.

Finally, since chopper cells are hypothesised to be important in speech processing due to their amplitude modulation tuning properties, we consider how deafferentation might affect this tuning in Figure 10 (with some caveats, see below). In this model, we modulate the auditory nerve fibre firing rate over time. In the normal cells with *N* = 50 auditory nerve fibre inputs, there is a peak in vector strength at a characteristic best modulation frequency (BMF). To model deafferentation, we reduce the number of input cells to *N* =10 and increase the synaptic weights *w* to restore the original mean input current *μ* to the chopper cell, as before. We found that for all parameter combinations we tried, deafferentation always had the effect of slightly reducing the vector strength at all modulation frequencies, and almost eliminating the peak.

**Figure 10:**
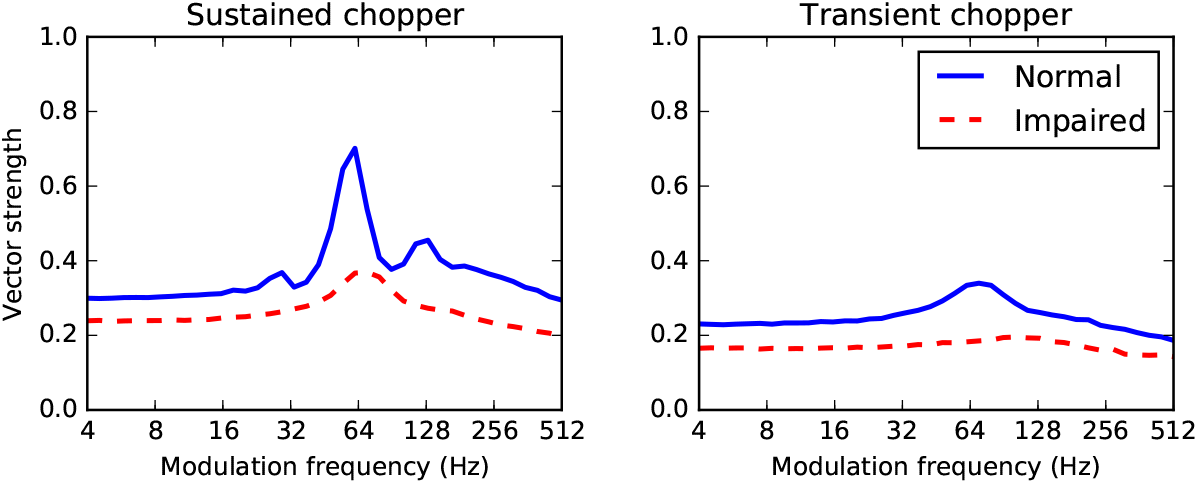
Effect of deafferentation on the temporal modulation transfer function for a simulated sustained (left) and transient (right) chopper cell in response to amplitude modulated tones. The normal cell vector strength at different modulation frequencies is shown with a solid blue line, and the impaired cell with a dashed red line.

Some caution is advised in interpreting these results. With an amplitude modulated tone signal, the assumptions in 2.2 will not hold. Specifically, the cell will not be in a stationary state and therefore we cannot ignore adaptation, and the amplitude varies so we cannot ignore compression. At high modulation frequencies, for example, the auditory nerve already shows a strong falloff in vector strength, probably because at high modulation frequencies the sidebands are outside the filter bandwidth (Joris et al., 2004), and this is not present in our model. For low modulation rates *f_m_* and small modulation depths m, however, these assumptions are not strongly violated and therefore these results may be suggestive. The normal cell responses are qualitatively similar to the experimental data, except at high modulation frequencies where we do not see the strong falloff in vector strength present in the data (Sayles et al., 2013). Note that we do not attempt to address the full range of mechanisms suggested to underlie chopper cell responses to AM tones. The sustained chopper model here with no inhibition is similar to Hewitt et al. (1992) and Lorenzi et al. (1995), and indeed Figure 10 is similar to Figure 14 of Hewitt et al. (1992) who also vary *N* while keeping the mean current fixed. The transient chopper model we use includes inhibition, but not the delayed inhibition with a different time course used in Nelson and Carney (2004).

## 4 Discussion

We can summarise the results above as follows: in a reduced mathematical model of the ongoing response of chopper cells, we can robustly reproduce sustained and transient chopper cell behaviour through a variety of mechanisms. This supports the idea that there may not be a single factor or mechanism governing chopping behaviour. However, in all cases we find that removing auditory nerve fibre inputs to the cell – for example as a result of noise-induced hearing loss (Kujawa and Liberman, 2009) – reduces the regularity of firing, which we would expect to have significant effects on subsequent processing in the auditory system. We now turn to a more detailed discussion of each of these points.

### 4.1 Reduced mathematical model

We chose to use an extremely simplified mathematical model of chopper cells in the style of Molnar and Pfeiffer (1968). This has several advantages and disadvantages. Firstly, the model makes relatively few assumptions about the nature of chopper cells, and therefore the results are rather general. Specifically, the same results would be obtained for a large class of more detailed models of chopper cells, and so our results also apply to these models. Secondly, the model has very few free parameters (just *μ, σ, τ, t*_ref_) and so it was possible to thoroughly explore the full parameter space of the model and show that the results are robust and do not require careful tuning.

However, the simplicity of the model comes with some limitations. The most important of these is that it can only predict the ongoing or stationary behaviour of the cell in response to a pure tone stimulus. This is unfortunate in two ways. First, the difference between sustained and transient chopper cells is partly defined by the time course of the regularity of firing: transient choppers are initially regular and rapidly become irregular. The model cannot reproduce this time course, however it is not too problematic, as the difference between sustained and transient choppers is highly correlated with their ongoing regularity (Blackburn and Sachs, 1989). Second, one of the most interesting aspects of chopper cell behaviour is their response to amplitude modulated sounds as captured by their modulation transfer function (e.g. Frisina et al. 1990; Sayles et al. 2013). Amplitude modulated stimuli violate the strict assumptions of this model, however for low modulation frequencies and depths the model produces results that are comparable with previous models (Hewitt et al., 1992; Lorenzi et al., 1995). This only covers one possible mechanism for reproducing modulation transfer functions of chopper cells. In future work, it may be possible to build a model that is only slightly more complex that can incorporate a wide range of possible mechanisms.

Finally, although the model is quite general, it does not cover all possible relevant factors. Synapses are assumed to be very fast acting, refractoriness is assumed to be only absolute and not relative, and so forth. These details can be included in the model, but doing do does not appear to alter the conclusions (results not shown).

### 4.2 Chopper cell mechanisms

We have shown that a wide range of factors can affect the firing regularity of chopper cells. Specifically, any factor that can robustly modify the value of *σ/μ* in equation 2 can generate both sustained and transient chopping. This shows that there does not need to be just one mechanism responsible for the difference between sustained and transient choppers, but does not rule out the possibility that in reality all chopper cells share a single mechanism for this.

On a similar theme, we can discuss whether cells ought to be divided into discrete or continuous classes. Traditionally, discrete classifications were used, so cells were defined as being choppers or primary-like, etc., and choppers were further subdivided into transient or sustained. However, Rothman and Manis (2003a,b,c) showed that there is a continuous distribution of the expression of various potassium currents in VCN cells that could lead to different behaviours. Typlt et al. (2012) carried out a clustering analysis of a large number of VCN cells and showed that there were no clear boundaries to categorise them into discrete groups. Our model provides further evidence to support this. Given that fundamental cell properties are varying on a continuous distribution, and that *σ/μ* will vary as a function of these properties, it is likely that the firing regularity will also vary in a continuous manner. This is further supported by our new analysis of the level-dependence of firing regularity, which shows that the large majority of cells change the regularity of their firing at different sound levels, and that in some cases this can cause them to cross the boundary between sustained and transient chopping. In conclusion then, we propose that it is better to describe specific behaviours in a continuous manner (e.g. more or less sustained/transient chopping).

This ties in with a general picture of neural functioning that has been developing in recent years, in which heterogeneity of cell properties is important for carrying out robust computations (Brette, 2012; Day and Delgutte, 2013; Garden et al., 2008; Goodman et al., 2013; Marsat and Maler, 2010; Padmanabhan and Urban, 2010; Raman et al., 2010; Zohar et al., 2013). In order to make further progress in our understanding of the function of the VCN, we may need to develop a clearer idea of how this heterogeneity may support downstream auditory computations. In particular, this may involve overlapping computations rather than a direct one-to-one correspondence between behaviour types and computations. This is well established in *C. elegans* (Bargmann and Horvitz, 1991; Biron et al., 2008; Clark et al., 2006), but has also been observed more widely (Rishel et al., 2013). For example, the cell properties that lead to chopping behaviour may contribute to several different auditory computations. This could partly explain why different mechanisms may be implicated in chopping behaviour.

### 4.3 Auditory processing and hearing loss

We identified mechanisms that may be associated with “hidden hearing loss”, similar to those proposed by Schaette and McAlpine (2011) (but without the distinction between low and high spontaneous rate fibres). Since regular firing is a defining feature of these cells, its disruption would indicate abnormal processing of auditory information both in the cochlear nucleus and downstream. For example, in Figure 10 we showed that after deafferentation chopper cells may have reduced tuning to amplitude modulation, which is likely to alter the representation of speech at the level of the inferior colliculus (Carney et al., 2015). Although we have shown this only for our chopper cells in our model, the same mechanism is likely to apply more widely in the cochlear nucleus and elsewhere in the auditory system. If individual spikes are stochastic, and there are fewer of them, then overall less information is being transmitted and the system as a whole would be expected to be noisier. This general hypothesis has been tested in a different way by Lopez-Poveda and Barrios (2013) and Lopez-Poveda (2014), who modelled the effect as a stochastic undersamplingofthe sound waveform. They found that simulating this stochastic undersampling impaired the complex sound processing more in noise than in quiet. Our results are consistent with this view of “hidden hearing loss”, in which deafferentation is associated with an increase in neural noise (Faisal et al., 2008) leading to a specific disruption to temporal processing. Finally, chopper units have been argued to play a role in level-dependent selective listening; that is they maintain a rate-place representation of complex sounds over a wide range of sound levels by responding to low threshold fibres at low sound levels and high threshold auditory nerve fibres at high sound levels (e.g. Lai et al. 1994). Interestingly it has been shown that it is the high threshold auditory nerve fibres that are preferentially lost following acoustic noise trauma (Furman et al., 2013), but we have not tested this idea explicitly.

In future work, it would be interesting to test these ideas both in more detailed models and experimentally. Our focus in this study was to demonstrate that this effect is robust and arises for a wide range of parameters in a reduced model. However, it would also be interesting to demonstrate the effect in a more detailed functional network model, to investigate in detail how deafferentation disrupts temporal processing. To verify experimentally whether CV is indeed affected by deafferentation consecutive to acoustic trauma, as predicted by our model, is difficult. It would ideally involve recording an individual cell before and long after an acoustic trauma, which is very challenging technically. More realistically we could hope to observe an overall reduction in the number of sustained choppers compared to transient choppers, although this would require recordings from a rather large number of cells in a large number of animals.

### 4.4 Reproducibility and robustness

We have made the code and an interactive version of this paper online at https://github.com/neural-reckoning/vcn_regularity. This has several aims. Firstly, we wish to allow readers to explore the consequences of changes in the parameters. A fixed set of figures in a printed paper only allows us to show a tiny snapshot of the complete parameter space, but with the interactive version the complete space can be easily explored without having to install any software locally. Secondly, we hope that it gives confidence in the results as it can be easily verified that the results presented here do not depend on highly tuned sets of parameters. Thirdly, by making the complete code available and possible to modify and run without any burden of installing complex software locally, we hope to encourage other researchers to use and build on this model with the minimum of effort. We would strongly encourage other authors to use the same or similar approach in the future, as the technology to do is now openly available and easy to use (Freeman et al., 2015).

## Acknowledgements

D.G., A.d.C., and C.L. were supported by two grants from ANR (HEARFIN and HEART projects). This work was also supported by ANR-11-0001-02 PSL* and ANR-10-LABX-0087.

